# Glycyrrhizin effectively neutralizes SARS-CoV-2 in vitro by inhibiting the viral main protease

**DOI:** 10.1101/2020.12.18.423104

**Authors:** L. van de Sand, M. Bormann, M. Alt, L. Schipper, C.S. Heilingloh, D. Todt, U. Dittmer, C. Elsner, O. Witzke, A. Krawczyk

**Affiliations:** Department of Infectious Diseases, West German Centre of Infectious Diseases, Universitätsmedizin Essen, University Duisburg-Essen, Germany; Department of Molecular and Medical Virology, Faculty of Medicine, Ruhr University Bochum, Bochum, Germany; Institute for Virology, University Hospital Essen, University of Duisburg-Essen, Essen, Germany

## Abstract

The newly emerged coronavirus, which was designated as severe acute respiratory syndrome coronavirus 2 (SARS-CoV-2) is the causative agent of the COVID-19 disease. High effective and well-tolerated medication for hospitalized and non-hospitalized patients is urgently needed. Traditional herbal medicine substances were discussed as promising candidates for the complementary treatment of viral diseases and recently suggested for the treatment of COVID-19. In the present study, we investigated aqueous licorice root extract for its neutralizing activity against SARS-CoV-2 *in vitro*, identified the active compound glycyrrhizin and uncovered the respective mechanism of viral neutralization. We demonstrated that glycyrrhizin, the primary active ingredient of the licorice root, potently neutralizes SARS-CoV-2 by inhibiting the viral main protease. Our experiments highlight glycyrrhizin as a potential antiviral compound that should be further investigated for the treatment of COVID-19.

## Main Manuscript

The newly emerged coronavirus, which was designated as severe acute respiratory syndrome coronavirus 2 (SARS-CoV-2) is the causative agent of the COVID-19 disease. Even presymptomatic patients or patients with mild symptoms are able to infect other people. Highly effective and well-tolerated medication for hospitalized and non-hospitalized patients is urgently needed. Besides compounds that were initially approved for the treatment of other viral infections such as remdesivir^1^, traditional herbal medicine substances were discussed as promising candidates for the complementary treatment of viral diseases and recently suggested for the treatment of COVID-19.

In the present study, we investigated aqueous licorice root extract for its neutralizing activity against SARS-CoV-2 *in vitro*, identified the active compound glycyrrhizin and uncovered the respective mechanism of viral neutralization.

Dried licorice roots were brewed in PBS at a concentration of 8 mg/ml (w/v) and the fluid was subsequently sterile filtered to obtain an aqueous licorice root extract. The neutralization capacity of licorice root extract was determined in cell culture by endpoint dilution. For this purpose, serial dilutions of the licorice root extract (0.004 mg/ml – 4 mg/ml) were pre-incubated with 100 TCID_50_ of SARS-CoV-2 for 1 hour at 37 °C and subsequently incubated on confluent Vero E6 cells grown in 96-well microtiter plates (pre-entry approach). After 48 hours, the cells were stained with crystal violet and analysed for plaque formation. Cytotoxicity was determined at four distinct time points (5 minutes, 12 hours, 24 hours and 4 hours) by using the “Orangu cell counting solution” (Cell guidance systems, Cambridge, United Kingdom), which is a WTS-8 based assay using NAD(P)H concentration and dehydrogenase enzyme activity to detect the cell vitality. The aqueous licorice root extract showed neutralizing effects even at a subtoxic concentration of 2 mg/ml, (Figure 1A and B). This concentration is lower than the normal consuming dilution e.g. in tea (12.5 mg/ml). Although licorice root tea may represent a good candidate for complementary use, the identification and characterization of the active compound is of great importance for a potential clinical application.

**Figure 1:**
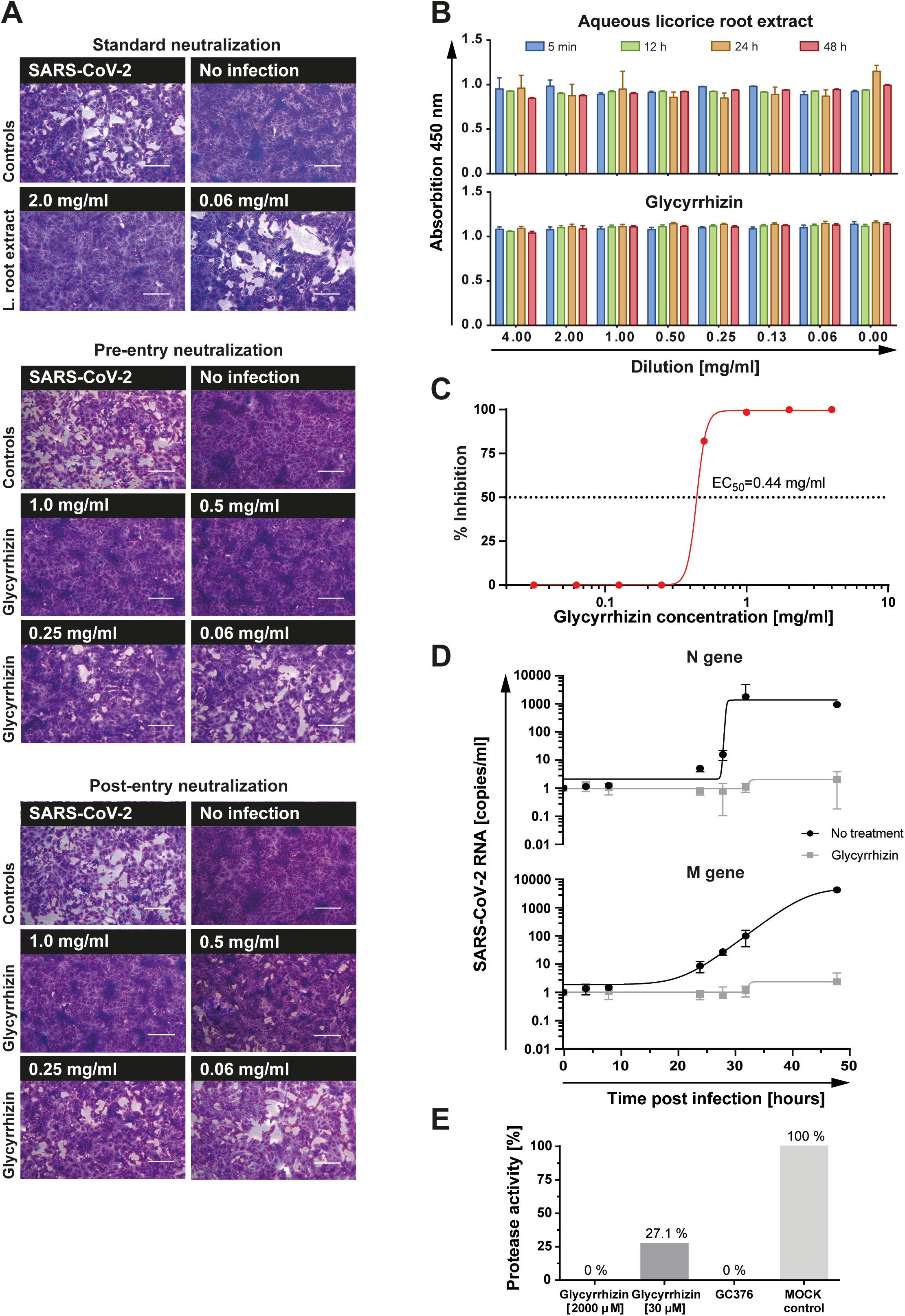
Antiviral efficacy of glycyrrhizin on the replication of SARS-CoV-2 in vitro. **A** Decreasing aqueous licorice root extract dilutions (0.004 mg/ml – 4 mg/ml) were pre-incubated with 100 TCID_50_/ml SARS-CoV-2 for 1 hour at 37°C and applied to a confluent layer of Vero E6 cells. After 48 hours of incubation, cell cultures were stained with crystal violet and analysed for plaque formation. The antiviral activity against SARS-CoV-2 was analysed under pre- and post-entry conditions. Descending glycyrrhizin concentrations (0.002 – 4 mg/ml) were pre-incubated with 100 TCID_50_ SARS-CoV-2 for 1 hour at 37°C (pre-entry condition) and subsequently added to confluent Vero E6 cells in 96-well microtiter plates for 48 hours. In a second approach, Vero E6 cells were inoculated with 100 TCID_50_ SARS-CoV-2 for 4 hours before the glycyrrhizin-containing inoculation medium with various glycyrrhizin concentrations (0.002 – 4 mg/ml end-concentration) was added (post-entry conditions). Plaque formation was evaluated after 48 hours post infection (p.i.). Bars represent 200 µm. **B** The toxicity of the treatment was tested by using “Orangu cell counting solution”. Different concentrations were incubated with a confluent layer of Vero E6 cells and evaluated at four time points (5 minutes, 12 hours, 24 hours, 48 hours). **C** Vero E6 cells were infected with 1000 TCID_50_/ 1.5 ml in different glycyrrhizin concentrations for 48 hours. The supernatant was titrated on microtiter plates in 1:10 dilutions to determine the viral loads in triplicates. The EC_50_ value was calculated using GraphPad Prism 8.0.1 (Graph Pad Software, San Diego, USA). **D** SARS CoV-2 RNA levels in supernatants of SARS-CoV-2 infected Vero E6 cells (500 TCID_50_) treated with 1 mg/ml glycyrrhizin or mock treated were determined at seven time points (0, 4, 8, 24, 28, 32 and 48 hours) p.i. by quantive RT-qPCR. **E** The inhibition of SARS-CoV-2 M^pro^ by glycyrrhizin was measured by using the “3CL Protease, MBP-tagged (SARS-CoV-2) Assay Kit” (BPS Bioscience, San Diego, United States).

Glycyrrhizic acid is a triterpene saponin and found in high concentrations in the root of the Glycyrrhiza glabra plant. It was described as an antiviral active ingredient of the licorice root and exhibits antiviral activity against herpes simplex viruses^2^, the human immunodeficiency virus as well as human and animal coronaviruses^3^. Lastly, an *in-silico* simulation study proposed an antiviral activity of glycyrrhizin against SARS-CoV-2, but this hypothesis remains experimentally unproved by now^4^. Based on our results with the aqueous licorice root extract, we investigated the antiviral activity of glycyrrhizin acid against a clinical SARS-CoV-2 isolate and subsequently examined the underlying mechanism of viral neutralization.

The neutralizing activity of glycyrrhizin against a clinical SARS-CoV-2 isolate was investigated in cell culture. Here, glycyrrhizin acid ammonium-nitrate was dissolved in DMEM containing 2% (v/v) FCS and 1% penicillin–streptomycin at 37 °C and adjusted to pH 7. A potential cytotoxic effect of glycyrrhizin was investigated as described above. No cytotoxic effect could be observed even at a concentration of 4 mg/ml (Figure 1B). The neutralization capacity of glycyrrhizin was determined by endpoint dilution. The antiviral activity against SARS-CoV-2 was analysed under pre- and post-entry conditions. Descending glycyrrhizin concentrations (0.002 – 4 mg/ml) were pre-incubated with 100 TCID_50_ SARS-CoV-2 for 1 hour at 37°C (pre-entry condition) and subsequently added to confluent Vero E6 cells in 96-well microtiter plates for 48 hours. In a second approach, Vero E6 cells were inoculated with 100 TCID_50_ SARS-CoV-2 for 4 hours before the glycyrrhizin-containing inoculation medium with various glycyrrhizin concentrations (0.002 – 4 mg/ml end-concentration) was added (post-entry conditions). Complete virus neutralization was achieved at subtoxic concentrations of 0.5 mg/ml under pre- and 1 mg/ml under post-entry conditions (Figure 1A and B). To further investigate the antiviral efficacy of glycyrrhizin, we determined the half-maximal effective concentration (EC_50_) sufficient to neutralize the virus. Confluent Vero E6 cells grown in 6-well plates were infected with 1000 TCID_50_ SARS-CoV-2 and at the same time treated with various concentrations of glycyrrhizin ranging from 0.0625 to 4 mg/ml. After 48 hours of incubation, the supernatants were harvested and the viral loads were determined by endpoint dilution. The experiment was performed in triplicates. The EC_50_ was calculated with 0.44 mg/ml, uncovering glycyrrhizin as a potent compound effective against SARS-CoV-2 (Figure 1C). The initial finding was supported by quantifying the SARS-CoV-2 RNA from the supernatants of SARS-CoV-2 infected cells treated with glycyrrhizin. Confluent Vero E6 cells grown in 24-well plates were infected with 500 TCID_50_ and simultaneously treated with 1 mg/ml of glycyrrhizin. Supernatants were collected at seven different time points (0 hours, 4 hours, 8 hours, 24 hours, 28 hours, 32 hours and 48 hours) post-infection. Viral RNA was purified from the supernatants with the “High Pure Viral RNA Kit” (Roche Diagnostics) and the genomic SARS-CoV-2 RNA was quantified by RT-qPCR. Therefore, primer targeting the viral M or N gene were used. M and N gene copy numbers were assessed using a 1:10 plasmid dilution series as reference (details and sequence information available upon request). Glycyrrhizin treatment significantly reduced the genomic SARS-CoV-2 RNA levels (Figure 1D). Taken together, we demonstrated that glycyrrhizin exhibited a high antiviral activity against SARS-CoV-2. Next, we investigated the underlying mechanism how glycyrrhizin may interfere with the virus replication. Recently, protease inhibitory activity of glycyrrhizin was predicted by *in silico* simulations^5^. The human transmembrane serine protease (TMPRSS2) was shown to cleave the SARS-CoV-2 Spike protein thereby facilitating the virus entry into the host cell^6^. However, since there was only a slight difference in antiviral activity of glycyrrhizin between pre- and post-entry conditions, and only a minor affinity was simulated for the interaction between glycyrrhizin and TMPRSS2, we concluded that glycyrrhizin neutralizes the virus by a mechanism different from inhibiting TMPRSS2. Thus, we focused on the SARS-CoV-2 main protease (M^pro^) as a potential target for glycyrrhizin^7^. M^pro^ is essential for processing the viral polyproteins that are translated from the viral RNA and thus, for virus replication^7^. Glycyrrhizin was suggested as a possible inhibitor of M^pro^ by *in silico* analysis, but this hypothesis has never been experimentally proven^5^. Here we provide evidence that glycyrrhizin potently inhibits M^pro^ activity *in vitro*. The inhibition of SARS-CoV-2 M^pro^ by glycyrrhizin was measured by using the “3CL Protease, MBP-tagged (SARS-CoV-2) Assay Kit”. Briefly, 90 ng of recombinant M^pro^ were incubated with two different concentrations of glycyrrhizin (30 µM and 2000 µM, dissolved in water). As control, the protease inhibitor GC376 was used. The enzyme-sample solution was incubated at room temperature for 30 minutes. The enzyme activity was measured at 360 nm excitation and 460 nm emission after overnight incubation of the inhibitor-M^pro^ mixtures with substrate (Dabcyl-KTSAVLQ↓SGFRKM-E(Edans)-NH_2_) at room temperature. Glycyrrhizin completely inhibited M^pro^ activity at a concentration of 2000 µM (1.6 mg/ml) and reduced its activity by 70.3% at a concentration of 30 µM (0.024 mg/ml).

Glycyrrhizin was clinically evaluated in the context of a clinical trial and described to be a safe and well-tolerated compound^8^. The pharmacological effects include antioxidative and anti-inflammatory, corticosteroid-like activities^9^. The potent antiviral activity as well as anti-inflammatory properties highlight glycyrrhizin as an excellent candidate for further clinical investigations in COVID-19 treatment. A case report described compassionate use of glycyrrhizin among other potential antivirals for the treatment of COVID-19^10^. Although the patient recovered from disease, further controlled studies are needed to prove the therapeutic effects of glycyrrhizin in COVID-19.

Taken together, we demonstrated that glycyrrhizin, the primary active ingredient of the licorice root, potently neutralizes SARS-CoV-2 by inhibiting the viral main protease. Our experiments highlight glycyrrhizin as a potential antiviral compound that should be further investigated for the treatment of COVID-19.

## ACKNOWLEDGEMENTS

This study was supported by the Stiftung Universitätsmedizin Essen (awarded to A. Krawczyk) and the Rudolf Ackermann Foundation (awarded to O. Witzke). The authors thank Barbara Bleekmann for excellent technical assistance.

## Address for correspondence

Adalbert Krawczyk, Department of Infectious Diseases, West German Centre of Infectious Diseases, Universitätsmedizin Essen, University Duisburg-Essen, 45147 Essen, Germany;

## AUTHOR CONTRIBUTIONS

L.V., A.K., M.A. and C.E. conceived and designed the experiments. L.V., M.A., M.B., L.S. and D.T. participated in multiple experiments; L.V., M.A. and A.K. analysed the data. L.V., A.K., C.E. and C.H. wrote the manuscript. A.K., O.W., M.A., L.V. and U.D. provided the final approval of the manuscript.

## References

1 Wang M, Cao R, Zhang L et al. Remdesivir and chloroquine effectively inhibit the recently emerged novel coronavirus (2019-nCoV) in vitro. Cell research 2020; 30:269–271.

2 Huang W, Chen X, Li Q et al. Inhibition of intercellular adhesion in herpex simplex virus infection by glycyrrhizin. Cell biochemistry and biophysics 2012; 62:137–140.

3 Cinatl J, Morgenstern B, Bauer G, Chandra P, Rabenau H, Doerr HW. Glycyrrhizin, an active component of liquorice roots, and replication of SARS-associated coronavirus. Lancet 2003; 361:2045–2046.

4 Chrzanowski J, Chrzanowska A, Graboń W. Glycyrrhizin: An old weapon against a novel coronavirus. Phytother Res 2020.

5 Srivastava V, Yadav A, Sarkar P. Molecular Docking and ADMET Study of Bioactive Compounds of Glycyrrhiza glabra Against Main Protease of SARS-CoV2. Materials today Proceedings 2020.

6 Hoffmann M, Kleine-Weber H, Schroeder S et al. SARS-CoV-2 Cell Entry Depends on ACE2 and TMPRSS2 and Is Blocked by a Clinically Proven Protease Inhibitor. Cell 2020; 181:271–280.e278.

7 Zhang L, Lin D, Sun X et al. Crystal structure of SARS-CoV-2 main protease provides a basis for design of improved α-ketoamide inhibitors. Science 2020; 368:409–412.

8 van Gelderen CE, Bijlsma JA, van Dokkum W, Savelkoul TJ. Glycyrrhizic acid: the assessment of a no effect level. Hum Exp Toxicol 2000; 19:434–439.

9 Kwon YJ, Son DH, Chung TH, Lee YJ. A Review of the Pharmacological Efficacy and Safety of Licorice Root from Corroborative Clinical Trial Findings. J Med Food 2020; 23:12–20.

10 Ding H, Deng W, Ding L, Ye X, Yin S, Huang W. Glycyrrhetinic acid and its derivatives as potential alternative medicine to relieve symptoms in nonhospitalized COVID-19 patients. J Med Virol 2020; 92:2200–2204.

